# Investigating Neurochemistry, Connectivity, and Audio Stimuli Relationship Among Surface and Depth Cortical Neurons

**DOI:** 10.64898/2026.03.16.711728

**Authors:** Nasim Winchester Vahidi, Sam Kassegne

## Abstract

In this study, we investigate the simultaneous recording of electrical and chemical signals within both the cortical surface and deep regions of the brain. This is made possible through the utilization of an innovative carbon based three-dimensional multi-functional neural probe. Our primary objectives are to explore in-depth the mechanisms of signal propagation among neuronal cells particularly within a three-dimensional framework and demonstrate initial progress in elucidating the interplay between electrical and chemical signals and their responsiveness to external stimuli variability.

**Approach:** Our innovative probe integrates *epi-cortical* (surface) and *intra-cortical* (depth) microelectrode arrays utilizing a two-dimensional thin-film microfabrication technique. This probe referred to as “*epi-intra*” has origami-like configuration and transforms from a two-dimensional structure into a three-dimensional configuration during implantation. Neural electrical signal recordings were conducted in the auditory region of an anesthetized European starling songbirds, whereas neurochemistry (dopamine) recordings were done simultaneously at Area X, with the animal subjected to conspecific songs as auditory stimuli.

**Main Results:** (i) This study introduces surface and depth neural recording in response to complex stimuli, such as bird songs using a three-dimensional probe with surface and depth microelectrodes. (ii) employing a *transfer entropy* model, a comprehensive connectivity map is established for neurons which are located on the surface of the brain, at a depth, or a combination of both, (iii) significantly, distinct spiking behavior in certain NCM (caudomedial nidopallium) neurons is observed during a specific phase of the stimulation coinciding with a peak in dopamine levels, which occurs with few milliseconds delay. This finding strongly indicates stimulus selectivity among specific neurons.

**Significance:** These findings demonstrate the creation of a connectivity map for neurons, whether located on the surface, at a depth, or a combination of both, derived from neuron recordings in response to a complex stimulus. Importantly, our study reveals that the strength of correlation and connectivity among neurons is most pronounced within surface neurons, followed by depth neurons, and comparatively weaker between surface and depth neurons. These discoveries hold significant promise in various applications, including the promise of large-scale neural electrical and electrochemical circuit mapping. Furthermore, they offer potential technology and therapeutic avenues for assisting, augmenting, or repairing human cognitive or sensory-motor functions.

## 1. Introduction

There is growing interest in elucidating the relationship between electrophysiology and neurochemistry within neural circuits through real-time recording of electrical and electrochemical signals in both the central and peripheral nervous systems. Understanding this interplay is critical for revealing neural communication at synapses [1-4], with important implications for developing treatments for neurological disorders affecting nearly 1 billion people worldwide [5]. In parallel, electrical stimulation is increasingly being explored as a therapeutic tool for conditions such as spinal cord injury, Parkinson’s disease, Alzheimer’s disease, and essential tremor [6-9]. Together, these advances highlight the need for a deeper understanding of how external stimuli interact with the brain’s electrical and electrochemical circuitry.

In the context of neural plasticity, the relationship among electrophysiology, stimuli, and neurotransmitters such as dopamine and serotonin has attracted substantial attention [10-12]. Although progress has been made in engineered plasticity for stroke and spinal cord injury (SCI), the links between neurochemistry and neuroplasticity, their underlying mechanisms, and whether they coincide with normalization of neurotransmitter chemistry remain incompletely understood [13-14]. To address this gap, we present a platform capable of simultaneous in vivo neurotransmitter detection and neural activity recording, enabling concurrent recording and stimulation of electrical and electrochemical signals and offering new insight into neurochemistry in neuroplasticity, particularly plasticity driven by audio stimuli [15]. In this study, we focus on dopamine (DA) because of its roles in motor activity, synaptic plasticity, reward processing [16], and regulation of blood flow and oxygen delivery in the frontal lobe [17]. Disruptions in DA signaling coordinated with electrical activity are implicated in disorders such as Parkinson’s disease and schizophrenia [18-19]. Using a songbird model, we examine DA release in the striatal song nucleus Area X, which depends on the social context of undirected versus directed singing [20]. Because Area X shares circuitry with HVC (high vocal center) and NCM (caudomedial nidopallium), we hypothesize that changes in electrical signaling within HVC and NCM alter DA concentrations in Area X [21]. This hypothesis forms a key basis of the present investigation.

Coupling these simultaneous recordings across a large neural circuit offers strong potential to advance our understanding of neural connectivity and electrochemistry in response to stimuli originating from either deep brain structures or the cortical surface [22]. Neurons rapidly propagate signals between deep and surface regions [12, 23-25], yet methods for studying these propagation mechanisms and the role of neurotransmitters in relation to stimuli remain limited [26]. To address this need, we perform simultaneous electrophysiological recordings from both the cortical surface and deep intracortical tissue using a 3-dimensional (3D) epicortical-intracortical configuration probe. This origami-like probe is enabled by our recently introduced pattern transfer technology, in which glassy carbon (GC) electrode structures are supported on polymeric substrates [27-30]. This microfabrication approach allows increasingly complex geometries, including 3D origami-style probes designed for both surface and depth recording [30]. Although fabricated in a 2-dimensional (2D) format, the device unfolds into a complex 3D structure. The resulting 3D neural probe integrates both surface (µECoG) and intracortical GC electrodes onto a single flexible thin-film platform, enabling recordings across a substantial tissue volume and offering a powerful tool for studying neuronal signal propagation and stimulus relationships. Our recent work demonstrated the effectiveness of this 3D origami-style “epi-intra probe” [30], which enabled us to characterize neural coding and stimulus reconstruction within a 3D brain volume [30-31]. This was achieved through analysis of both local field potentials (LFPs) and high-quality, stimulus-locked auditory single-unit responses from deep and surface regions [31].

In this study, we use origami-style 3D probes to investigate neural functional connectivity patterns within a 3D brain volume in response to audio stimuli using the transfer entropy (TE) model, focusing on neurons located at both the cortical surface and in deep brain regions [32-33]. The TE model, aligned with Granger causality principles, measures mutual information (MI) between the past states of a source signal X and the future state of a target signal Y while conditioning on the past of Y, thereby quantifying improvements in prediction using log-loss [34-36]. This framework provides insight into relationships between neural activity and audio stimuli across neurons recorded by our 3D probe configuration. Although functional connectivity has been widely studied, varying definitions and analytical methods continue to generate debate [37-38]. In this context, TE offers a promising and comprehensive approach for defining connections, estimating their strength, and determining the direction of information flow between neurons across regions. Here, connectivity direction refers to the dominant flow of information; a positive result indicates that information primarily flows from X to Y, or that one system provides greater predictive information about the other [39]. Overall, connectivity analysis in this study offers insight into the relationship between neural activity and audio stimuli across neurons recorded by the 3D probe and may also aid in understanding, predicting, and diagnosing neurological disorders, as reduced neuronal connectivity has been associated with Parkinson’s disease [40].

## 2. Materials and Methods

We utilized a 3-dimensional probe equipped with both surface and depth microelectrodes, as shown in Figure 1B. This probe, introduced in an earlier publication, features eight surface electrodes (50 µm diameter, 100 µm center-to-center pitch) and four depth glassy carbon electrodes (100 µm pitch) [31]. The microfabrication procedure is described in detail in **Figure S1** and our previous publications [28-31]. Briefly, SU-8 negative photoresist (Microchem, MA) was spin-coated on a silicon wafer and patterned by UV exposure (∼400 mJ/cm^2^), followed by post-baking and pyrolysis at 1000°C in an inert N_2_ environment as described previously [28-29]. A 5 µm layer of photo-patternable polyimide (HD 4100; HD Microsystems, DE, USA) was then spin-coated (2500 rpm), soft baked (90°C for 3 min, then 120°C for 3 min), and patterned (∼400 mJ/cm^2^ UV exposure), followed by partial curing at 300°C for 60 min under N_2_. Pt traces with a Ti adhesion layer were subsequently patterned by metal lift-off. For insulation, an additional 6 µm layer of polyimide HD 4100 was spin-coated (300 rpm), patterned (400 mJ/cm^2^), and cured (350°C for 90 min) under N_2_. A further 30 µm layer of polyimide (Durimide 7520, Fuji Film, Japan) was spin-coated (800 rpm, 45 s) and patterned (400 mJ/cm^2^) to reinforce the penetrating portion of the device. The microfabrication steps for the voltammetry probe used for neurotransmitter detection are shown in **Figure S2** [41-42].

**Figure 1.**
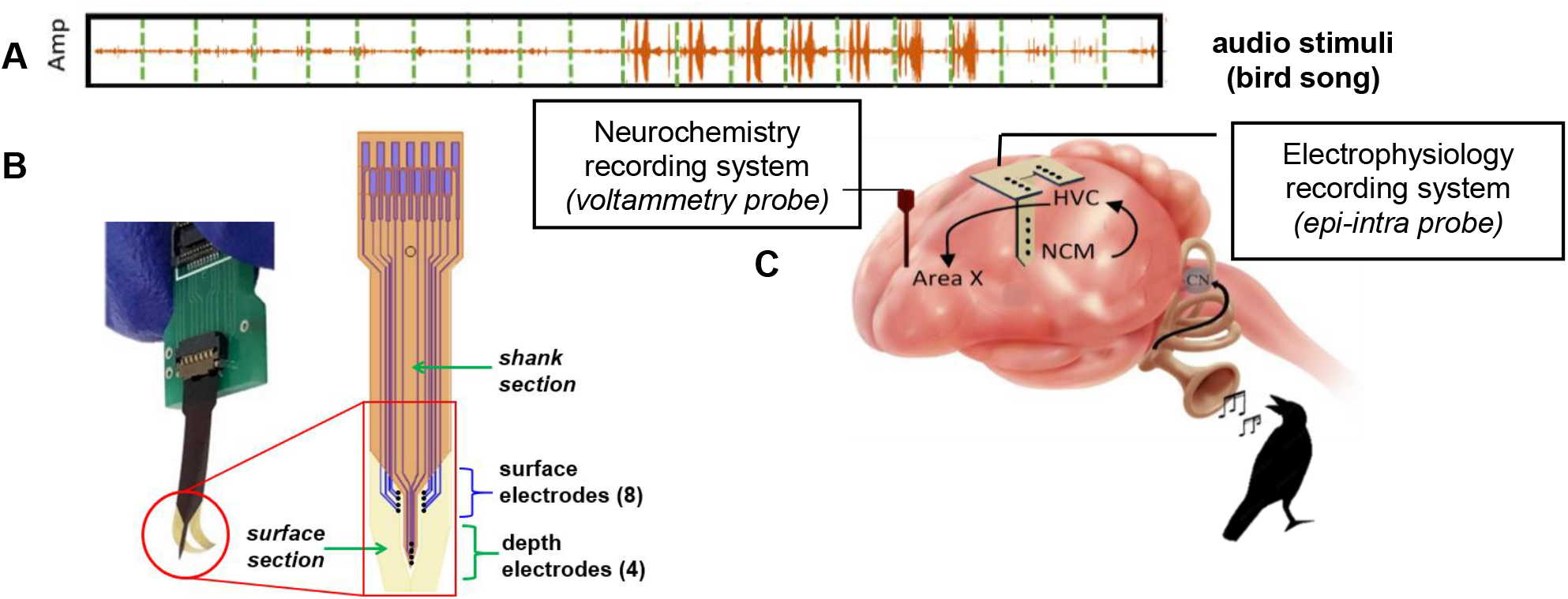
Schematic of 3D probe placement for surface and depth recording (A) audio stimulus (B) 2D schematic of epi-intra probe (4.15mm wide and 18.0mm long) (C) Schematic depicts the placement of the epi-intra and the voltammetry probes. The epi-cortical electrodes contact the HVC region, while the intra-cortical electrodes penetrate into the NCM area. Additionally, neurotransmitter data is recorded from Area X using voltammetry electrodes. The figure also illustrates the neural pathway among NCM, HVC, and Area X.

Experiments were performed on adult male European Starling songbirds under a protocol approved by the UCSD Institutional Animal Care and Use Committee (IACUC protocol number S05383). The flap portion of the probe, containing the surface electrodes, was placed on the high vocal center (HVC), while the shaft containing depth electrodes was inserted into the caudomedial nidopallium (NCM). The epi-intra probes enabled simultaneous recordings from the brain surface (HVC) and a deeper auditory region (NCM). Notably, this cortical region is analogous to the auditory association cortex of the mammalian temporal lobe [43]. In addition, the shaft of the voltammetry probe, containing four GC microelectrode sites, was inserted into Area X (Figure 1C), allowing us to record dopamine release in Area X, which is connected to HVC and NCM [21]. This study focuses on the relationship between neurochemical data from Area X and electrical signals recorded from neurons in HVC and NCM. To examine neural activity dynamics and adaptation to stimuli, we applied the TE model to 20 consecutive response segments from a total of 11 neural recordings (Figure 4). Optical images of the epi-intra and voltammetry probes are shown in **Figure S3**, schematics of the electrophysiology and neurochemistry recording setup are shown in **Figure S4**, and **Section S1** describes electrochemical characterization of the microelectrodes, including impedance spectroscopy data shown in **Figure S5**.

### 2.1. Audio Stimuli and Neural Response Recording

We conducted neural electrical signal recordings on three adult male European starling songbirds (n = 3), each with an average weight of 85 g. During the surgical procedures, the epi-intra electrode arrays were advanced into the auditory region of the anesthetized birds’ brains. The birds’ heads were positioned at a 22° angle using the intersection of the ear bars as the reference point. The probe was then placed at coordinates 500–1000 µm caudal and 500–1000 µm lateral on the right side of the Y-sinus. The shank containing the depth electrodes was lowered to a depth of 3000 µm until the surface array established close and conformal contact with the cortex. At these coordinates, the surface electrodes recorded neuronal signals from the HVC region, while the depth electrodes recorded signals from the NCM area.

After collecting both the stimuli and neural response data, we processed the recordings offline using MATLAB [44], Python [45], and Java platforms. Specifically, power spectrograms were generated for both the bird song stimuli and the corresponding 11 neural response channels using MATLAB. The spectrograms were computed with nfft = 128, a Hanning window of 128 samples, and 5% overlap. Following implantation of the epi-intra electrodes into the HVC and NCM regions, three recorded starling bird songs were played in random order and repeated 20 times. Simultaneously, neural responses from 11 channels were recorded and filtered within the frequency range of 10 Hz– 10 kHz, with a gain of 5K. Recordings were performed using A-M Systems (Sequim, WA) over a 90-minute session. The signals were subsequently digitized with an additional 10× gain using an A/D converter (CED Power 1401). **Figures S6** and **S7** present representative examples of the recorded neural data. Figure S6 shows high-pass filtered signals recorded from both surface and depth electrodes in response to a 20-second bird song stimulus. Figure S7 illustrates an example of spike detection and analysis from the NCM region, including the stimulus spectrogram, raster plots across repeated trials, averaged spike responses, extracted spike waveforms, and interspike interval distributions.

### 2.2. Neurochemistry Recording

To measure dopamine levels, we employed fast-scan cyclic voltammetry (FSCV) using a separate depth probe equipped with voltammetry microelectrode recording sites [41,42]. While, in principle, one of the channels of the *epi-intra* probe could be used for voltammetry, the use of a separate probe was motivated by two considerations: (i) the dopaminergic region Area X is located away from the HVC and NCM areas and (ii) separate hardware is required for FSCV with its own wiring. The probe with 30 μm x 50 μm size and 220 μm spacing voltammetry microelectrodes [42] was strategically placed in Area X of Starling bird brains, with the birds’ heads positioned at a 22o angle (with the intersection of ear bars as the reference point) and coordinates set at *3*.*9 AP, 1 ML, and -5 depth*. While playing bird song stimuli, we simultaneously recorded the concentration of dopamine using WaveNeuro® FSCV Potentiostat System (Pine Research, NC) in conjunction with brain electrical responses. For FSCV, a triangular waveform was scanned at *500 V/s* from *-0*.*5V to 1*.*3V* (vs. Ag/AgCl reference) and a frequency of 10 Hz. Analyte presence is detected by observing their specific oxidation and reduction peaks (Figure 2) [29,41-42].

**Figure 2.**
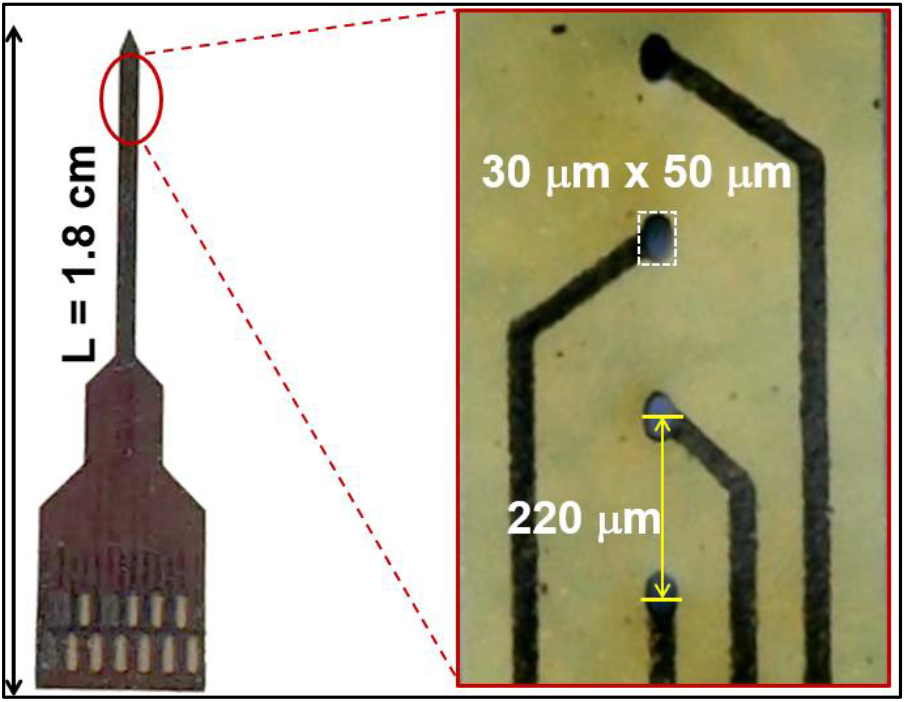
4-channel voltammetry probe for neurotransmitter detection. Each of the voltammetry electrodes are of 30 μm x 50 μm size and 220 μm spacing.

### 2.3 Functional Connectivity and Cross-Correlation Analysis

**Section S2** describes spike detection, sorting, and signal processing steps. We investigated *functional connectivity* (FC) between surface and depth neurons using the *transfer entropy* (TE) model. To compute the normalized functional connectivity across recorded sites, we utilized MATLAB in conjunction with *Java Information Dynamics Toolkit (JIDT)* and *ToolConnect* [46-47]. These platforms serve as an information-theoretic toolkit tailored for studying complex system dynamics. These toolkits provide a range of tools for quantifying transfer entropy, mutual information, conditional mutual information, and monitoring the spatiotemporal dynamics of these operations.

## 3. Results

### 3.1 Relationship Among Bird Song Stimuli, Brain Responses, Correlation, and Connectivity

We applied the TE model to analyze the responses of *11* neural channels, consisting of *7* surface channels (*epi* in red line) and 4 depth channels (*intra* in blue line) (Figure 3E). This analysis provided insights into time-series connectivity, time delay (indicating information flow direction), and correlations among these neural responses (Figure 3A-C) [48]. To visualize the changes in connectivity among neuronal responses to audio stimuli, we show the combined connectivity strengths and directions in circular graphs (Figure 3D) [49-50]. In Figure 3A, the color map represents the connectivity strength between each pair of electrodes/neural responses. A brighter yellow color indicates more robust connectivity, while a darker blue color indicates weaker connectivity. The color bar on the right side provides a visual representation of connectivity strength, ranging from +1 (strongest) to 0 (weakest), between two electrodes. Figure 3B demonstrates the matrix of time delay or direction of information flow among electrodes [51]. Figure 3C presents cross-correlations between surface (*epi*) and depth (*intra*) channels, with the color bar indicating the intensity of these cross-correlations. Figure 3D features a circular graph that combines the information from Figures 3A and 3B. In this figure, the thickness of the connecting lines reflects connectivity strength, with thicker lines indicating more robust connections. Additionally, the colors on these lines indicate the direction of information flow, transitioning from red to green. It is important to note that in cases of high connectivity strength between two electrodes, the algorithm may not determine direction, resulting in blue colors. Extended version of Figure 3 is given in **Figure S8**.

**Figure 3.**
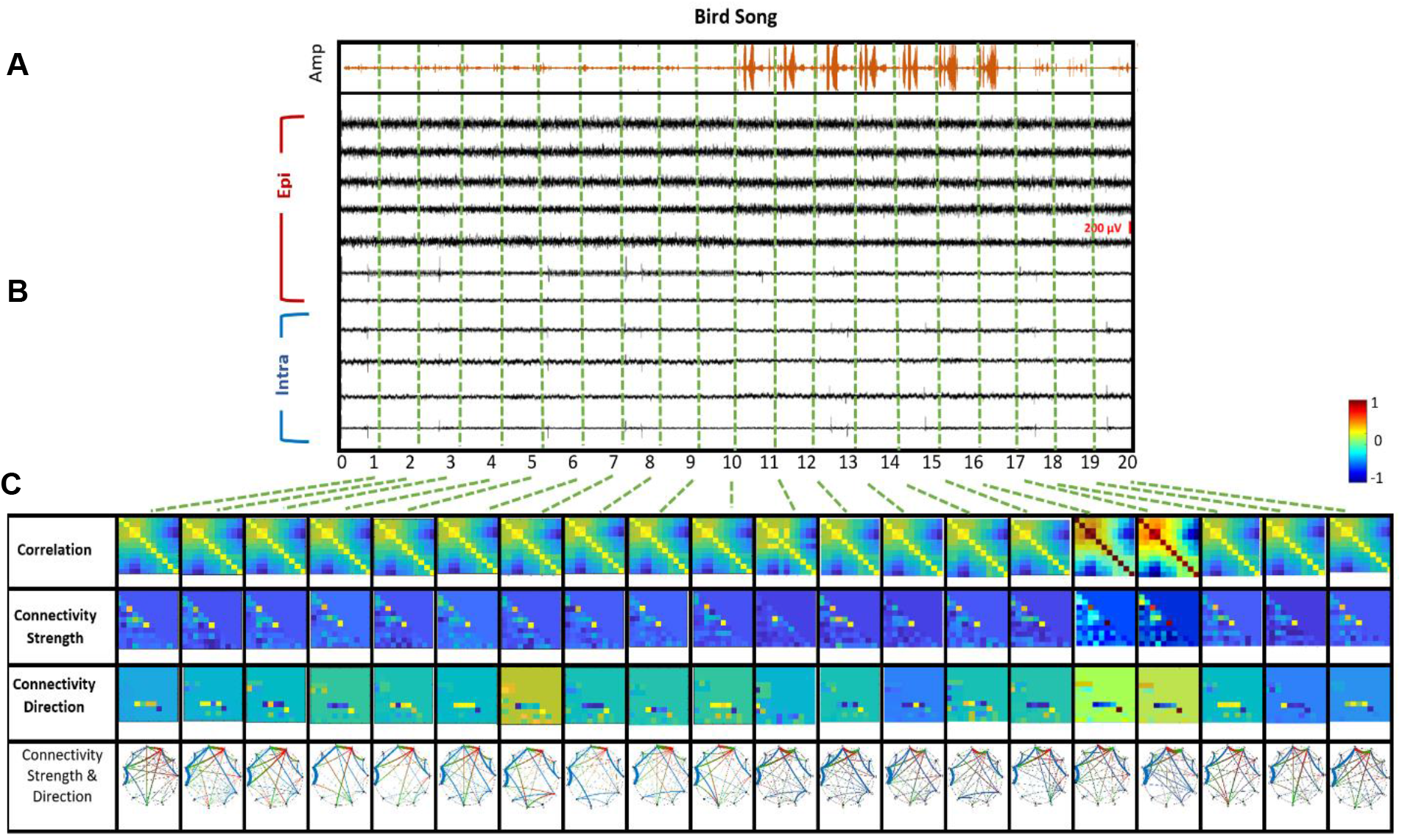
**(A)** Example of normalized functional connectivity among 11 neurons, with surface electrodes (epi) in red and depth electrodes (intra) in blue. The color bar represents the strength of connections between pairs of electrodes. (B) Time delay between these 11 neurons, with colors indicating the direction of information flow between each pair (C) Cross-correlation between surface and depth channels for 20 sec signals. Color bar indicates the intensity of correlation between these channels. (D) The circular graph combines both connection strength and direction of information flow among the 11 surface and depth neurons. Thicker lines indicate stronger connectivity, while thinner lines suggest weaker connections between each pair of electrodes. Color lines represent direction of information flow, from red to green. Blue indicates strong connectivity between two electrodes. (E) The surface (**red**)-depth (**blue**) combination electrode array with labeled channel numbers.

In Figure 4, we present an analysis of neural responses to 20 seconds of bird song stimuli. This graph offers a detailed view of the relationships between bird song stimuli, brain responses, correlation, functional connectivity, and time delays among the 11 electrodes on both the brain surface and depth. In the top row (Figure 4A), we observe the amplitude of the bird song stimulus over 20 seconds. Below this, the corresponding 20-second neural responses recorded from 11 electrodes, filtered within the range of 10 Hz to 10 kHz, are displayed. The seven channels recorded from the brain surface are highlighted in red brackets (*epi*), while the four channels from depth electrodes are marked in blue brackets (*intra*). To facilitate analysis, green vertical dotted lines divide the 20 seconds of connectivity data into one-second sections (Figure 4B).

**Figure 4.**
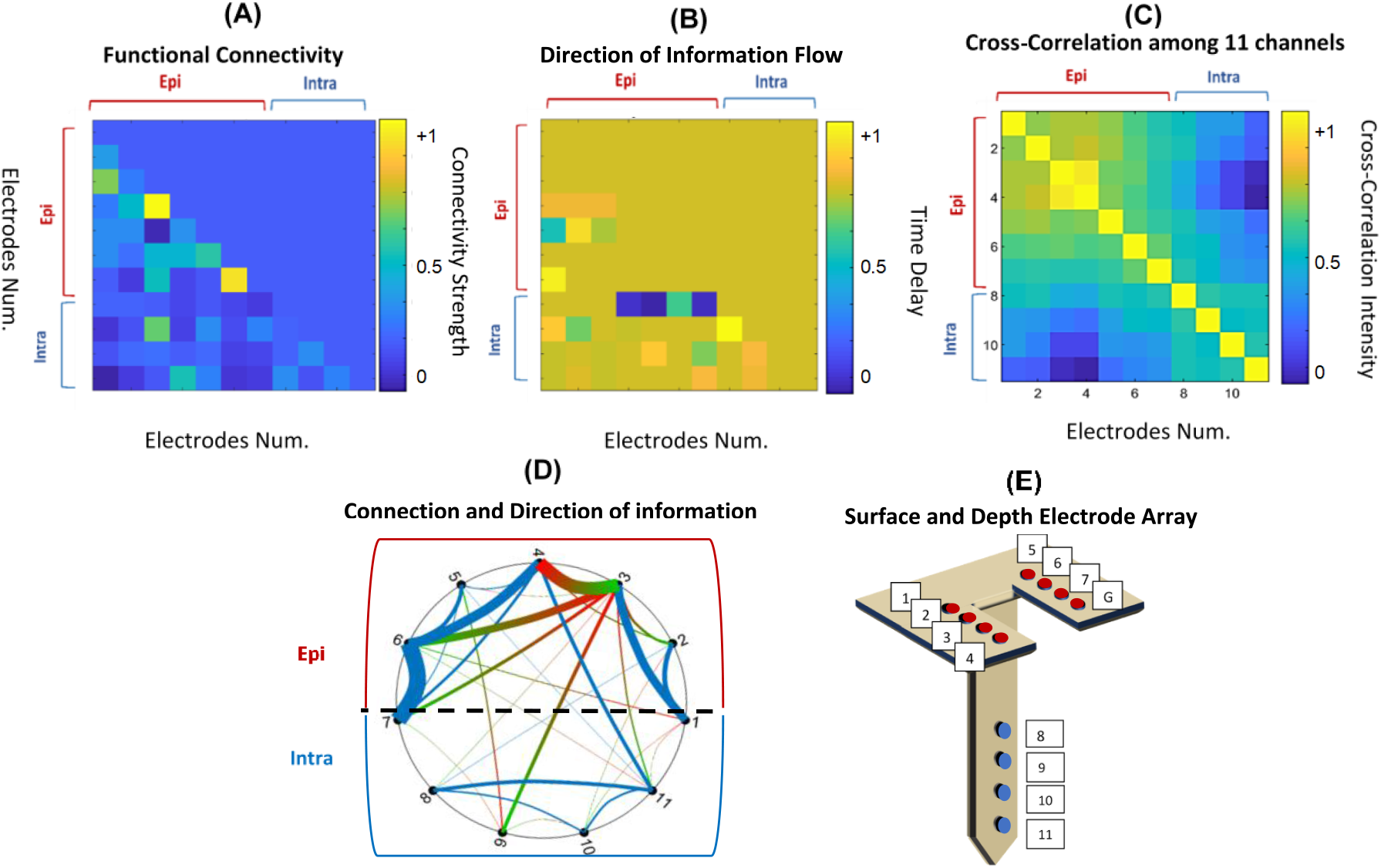
Relationship among bird song stimuli, brain responses, correlation, functional connectivity strength, and direction among 11 neurons in the brain surface and depth. (A) The first row shows the amplitude of a 20-second bird song stimulus. **(B)** Corresponding 20-second neural responses recorded from 11 electrodes are displayed. These recordings were filtered at 10Hz-10KHz. Seven channels recorded from the brain surface are shown in red brackets, and four channels recorded by depth electrodes are indicated by blue brackets. The green vertical dotted lines divide the 20 one-second concatenated neural response segments. **(C)** In the bottom graph, analyses are presented for the 11-channel brain responses during 20 consecutive one-second intervals (from the first to the fifth row). The first row displays correlations among 11 recorded electrodes from the brain surface and depth during 20 one-second neural responses. In the second row, the connectivity maps define the strength of connectivity between neurons on the brain surface and depth during the playback of stimuli. Third row shows the time delay map, defining the direction of information flow between each pair of neurons. The color bar specifies the strength of connectivity and direction of information flow between two neurons, ranging from +1 to -1. Lastly, in the fourth row, circular graphs combine information about connectivity strength and direction between each pair of neurons. The thickness of the connecting lines indicates connectivity strength, while colors represent the direction of information flow from red to green between two neurons. Blue color indicates the absence of a detected direction, which appears to occur between neighboring sites.

Figure 4C contains four rows, displaying correlation, connectivity, direction, and circular graphs among the 11-channel brain recordings for each one-second section. In the first row, the correlation matrix shows cross-correlations between responses from the surface and deep brain regions. In the second row, connectivity maps reveal the strength of connections between electrodes on the brain surface and depth during the playback of the 20-second bird song stimuli. The third row focuses on the time delay map, which defines the direction of information flow between pairs of electrodes. Finally, the fourth row presents circular graphs that combine information about connectivity strength and direction between each pair of electrodes. Overall, these circular graphs provide insights into changes in neuron connectivity, transient connections, and their strengths within the 3D volume of the brain, encompassing both surface and depth neurons during the bird song performance. This observation indicates that certain groups of neurons could be responsive to specific elements of the stimulus, aiming to interpret particular components of the stimuli [12, 53]. Interestingly, some electrodes maintain more permanent connections despite variations in bird song amplitude and content, as seen in the consistent connectivity between signal recordings at electrodes 3 & 4, 6 & 7, and 6 & 4 across all 20 circular graphs (refer to Figure 3D for a magnified *1-second* time frame from the analyzed 20 seconds).

### 3.2 Effects of Bird Song Stimulus Power and Frequency on Neural Connectivity Dynamics Among Surface and Deep Neurons

Twenty *1-second* segments of bird song data are given in Figure 5A, with Figure 5B illustrating the power (dB/Hz) versus frequency (Hz) relationship for these ten consecutive segments. The estimation is based on the *Welch* method (MATLAB’s *pwelch* function), which calculates the power spectral density (PSD) of the bird song. This analysis reveals that within the song’s frequency range spanning from *1 Hz - 20 KHz*, two dominant frequency ranges emerge: *1-300 Hz* (shown in blue) and *300-600 Hz* (shown in red). These specific stimulus frequencies may influence the connectivity and direction of information flow among neurons located on the cortical surface and those at depth [54-55].

**Figure 5.**
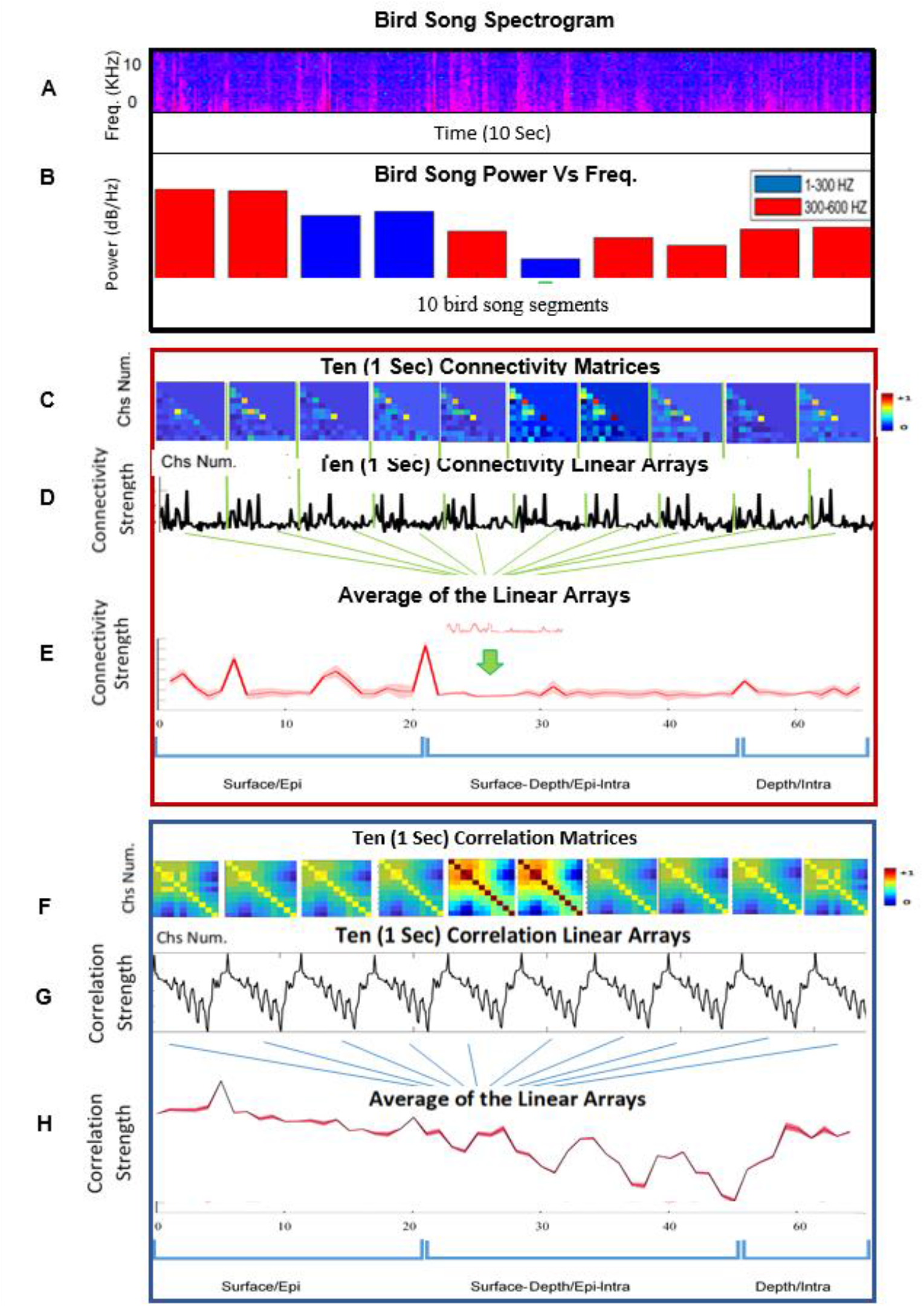
(A) Spectrogram of a 10-second bird song stimulus. (B) Power (dB/Hz) vs. frequency (Hz) plots for 10 consecutive bird song segments.(C) Ten 1 Second Connectivity Matrices (D) Conversion of the connectivity matrices into linear arrays, each containing three sections: epi, epi-intra, and intra. This process is repeated for the ten connectivity matrices. (E) The ten connectivity matrices converted to ten arrays of data, each containing three sections: epi, epi-intra, and intra. Next, the ten arrays are averaged. The pink band represents the standard deviation among the ten connectivity arrays, while the red line shows the mean for surface neurons, surface-depth neurons, and depth neurons in response to a ten-second stimulus. (F) Ten Connectivity Matrices. (G) Each correlation matrix is divided into three sections: epi, epi-intra, and intra. Next, each matrix is converted into a linear array. These arrays are then concatenated. This process is repeated for the ten correlation matrices. (H) The ten arrays are averaged. The pink band demonstrates the standard deviation among the ten correlation arrays, while the red line shows the mean.

### 3.3 Functional Connectivity Among Surface and Depth Neurons

In this section, Figures 5C-D explore the functional connectivity between surface and depth neurons. The connectivity matrices used in this analysis were derived from the last ten seconds depicted in Figure 4C. These matrices were categorized into three sections: *epi, epi-intra*, and *intra*, as illustrated in Figure 3A. Each of these sections was then transformed into three linear arrays. This transformation was repeated for the ten connectivity matrices, culminating in Figure 5D, which displays the converted matrices as ten data arrays. Figure 5E presents an average of these ten arrays. Within this graph, the pink band indicates the standard deviation observed among the ten connectivity arrays, while the red line represents the mean. The red mean line acts as an indicator of the strength of neural connections, and the standard deviation illustrates variations in neural connections among surface neurons, surface-depth neurons, and depth neurons in response to the ten seconds of stimuli. From this analysis, it is evident that connectivity is stronger among surface neurons compared to connections between surface and depth neurons [11].

### 3.4 Correlation Dynamics Among Surface and Depth Neurons

Here, we adopted an approach similar to that presented in Figure 5D to analyze the dynamics of correlation strength among surface and depth neurons. We utilized the final ten correlation matrices from the first row of Figure 3C, dividing each matrix into three sections: *epi, epi-intra*, and *intra*, akin to the depiction in Figure 2C. These three matrices were subsequently transformed into linear arrays and then concatenated. This process was repeated for all ten correlation matrices, as visualized in Figure 5F. Figure 5G illustrates the transformation of these matrices into ten data arrays, which were further averaged to generate the composite representation seen in Figure 5H. Within this depiction, the pink band represents the standard deviation across the ten correlation arrays, while the red line indicates the mean. The red line indicates the strength of neuronal correlation, demonstrating its highest levels among surface neurons, a decrease among surface-depth neurons, and a slight increase among depth neurons.

### 3.5 Relationship Between Neurochemistry, Neuronal Responses, and Stimuli

This study illustrates that changes in the electrical properties of the neuron population in HVC and NCM are temporally correlated with dopamine secretion in Area X. To investigate this relationship, we analyzed the temporal dynamics of neuronal firing, dopamine secretion, and the birdsong stimulus (**Figure 6**). **Figures 6A-B** showcase a 40 ms birdsong wave and its corresponding spectrogram. **Figure 6C** presents FSCV data displaying alterations in dopamine levels recorded from Area X. **Figure 6D** depicts the dopamine voltage-current graph, generated by averaging all current signals across voltage sweeps, illustrating the timing and location of oxidation and reduction events. Figure 6E shows dopamine current alterations over time, with an oxidation peak indicating that dopamine levels reached their maximum during a specific part of the stimulus. Lastly, **Figure 6E** illustrates a neural spike recorded by a depth electrode in NCM, with the dotted vertical line indicating the time difference between electrical and chemical signals.

**Figure 6.**
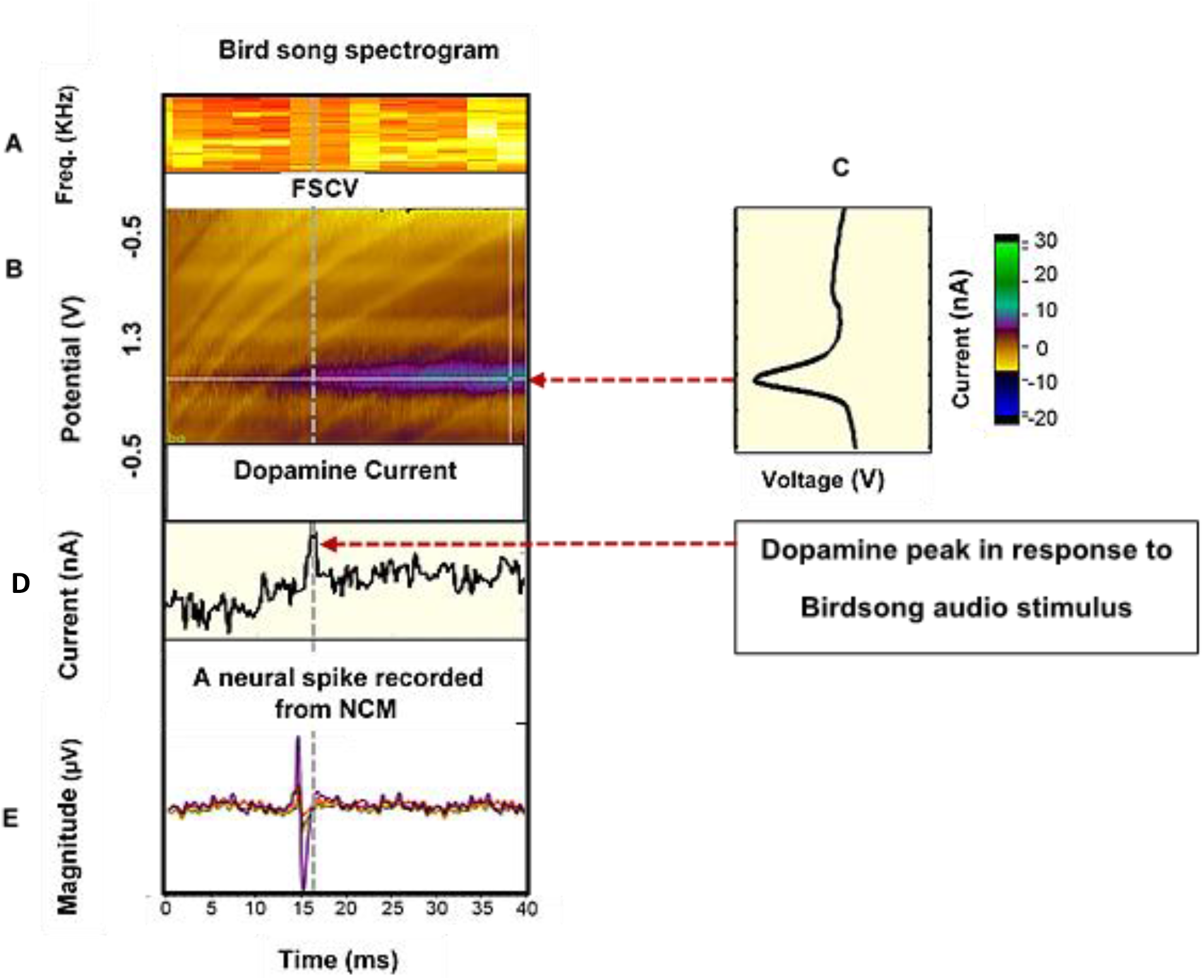
(A-B) Depicts a 40ms birdsong signal and its corresponding spectrogram. (C) Displays Fast-Scan Cyclic Voltammetry (FSCV) data capturing dopamine alterations recorded from Area X and the dopamine voltage-current graph, indicating where and when oxidation and reduction processes occurred. (D) Shows the oxidation current-time graph, highlighting when dopamine reached its maximum levels during a specific part of the birdsong stimulus. (E) Exhibits a neural spike recorded from a depth electrode in NCM, with the dotted vertical line denoting the time delay between electrical and chemical signals during the 40ms birdsong stimulus.

The observed temporal correlation between HVC and NCM neural firing and dopamine secretion in Area X is based on the time delay identified in **Figures 6C-D**. Specifically, the dotted vertical line in **Figure 6E** highlights the temporal alignment between neuronal activity and changes in dopamine current. While this provides suggestive evidence of a relationship, we recognize the need for more robust statistical support. To strengthen our claim, we performed the following statistical analyses: Pearson’s correlation coefficients were calculated between HVC firing rates and dopamine current values over time. The results revealed a statistically significant positive correlation (*p* < 0.05), supporting a temporal association between the two variables. To assess potential causality, a time-lagged cross-correlation analysis was performed. The analysis demonstrated that changes in HVC firing rates precede alterations in dopamine current, with a statistically significant peak correlation observed at the identified delay (**Figure S9 A-B**) The signals demonstrate a temporal correlation, with neural firing potentially influencing dopamine secretion. This observation aligns with the study’s claim that HVC neuronal activity and dopamine secretion in Area X are temporally linked, although causation cannot be definitively established. To ensure the temporal correlation was not due to chance, permutation tests were conducted by randomly shuffling the data 1,000 times. The observed correlation was significantly higher than the null distribution (p < 0.01). **(Figure S9 C**).

## 4. Discussion and Conclusion

In this study, we assessed the impact of stimulus variability on auditory neurons’ responses and neurochemistry using the *transfer entropy* model. The combination of tools employed here offers the potential to serve as a platform for advancing research in Brain-Computer Interfaces (BCI) [57-59]. We conducted this study by recording neuron responses in reaction to a complex stimulus, such as a bird song, using the 3D *epi-intra* probe. Subsequently, we generated a connectivity map for neurons on the brain’s surface, in deeper regions, or in combinations of thereof, utilizing the *transfer entropy* model, which leverages m*utual information* among neurons to predict their network dynamics and directions.

Additionally, our study employed circular graphs to shed light on various aspects of neuron connectivity. This includes electrodes connectivity strength and direction of information within the 3D brain volume, involving both surface and depth neurons during bird song performance. Furthermore, our analysis highlights that surface neurons demonstrate stronger connectivity and correlations with each other than depth neurons, primarily due to closer physical proximity that enables more direct interactions. These findings are underscored by connectivity maps and correlation matrices. Conversely, the strength of neuronal correlation and connectivity diminishes across surface-depth interactions and shows a slight resurgence among deep neurons, suggesting a complex pattern of neuronal interactions across different cortical layers. Our investigation also delved into the influence of bird song stimulus power and frequency on neural connectivity dynamics among surface and deep neurons. This analysis revealed that for songs within the frequency range of *1 Hz - 20 KHz*, two frequency ranges, *1-300 Hz* and *300-600 Hz*, exhibit dominance. These specific stimulus frequencies may impact connectivity and influence the direction of information exchange among cortical surface and deep cells.

In the context of exploring the intricate relationship between electrophysiology, neurochemistry (dopamine), and stimuli, our study investigated the chemical and electrical signals and their dynamics in response to stimulus variability. Our findings showed that during specific segments of the stimulus, certain neurons in the NCM exhibited spiking behavior, and dopamine levels peaked with an approximately *5ms* delay. This suggests potential stimulus selectivity by specific neurons, indicating their adaptability to the bird song stimulus. In other words, certain neural groups seem to respond selectively to specific elements within the stimulus, with the purpose of interpreting distinct components of the stimuli. In a previous work, we had shown successful *in vivo* measurement of spontaneous DA concentration in the Striatum of European Starling songbird through FSCV using a *4-channel* depth probe [41]; but with no simultaneous electrical signaling recorded. This current work which uses a combination of *12-channel* 3D neural response recording probe and *4-channel* voltammetry recording probe -- both with GC electrodes -- represents a significant progress where dopamine measurement is accompanied by a simultaneous electrophysiology recording. This result could be viewed in a larger context of providing an initial finding in large-scale simultaneous electrophysiology and neurochemistry recordings that will require both types of recordings at depth and potentially surface locations. Such comprehensive sets of data could then potentially enable the development of deeper understanding of the inter-relationship between these two modes of neural communication.

Our findings demonstrate that song stimuli induce neural discharges in the NCM area and delayed dopamine release in Area X. This observation aligns with the established role of Area X in processing song-related auditory and motor signals, particularly in songbirds. The delayed dopamine release may reflect modulatory input influencing reward prediction or motor learning associated with song production and perception. In addressing the neural pathways contributing to dopamine release in Area X, we recognize that dopaminergic projections to Area X predominantly originate from the ventral tegmental area (VTA), as previously described in the literature [62]. The HVC-to-Area X pathway is known to be primarily glutamatergic, conveying excitatory signals critical for song timing and motor control. Meanwhile, the functional interaction between HVC, NCM, and dopaminergic modulation in Area X remains an area of active investigation. While our data suggest an indirect relationship between HVC/NCM activity and dopamine release in Area X, it is possible that this is mediated by secondary circuits involving VTA. For example, song stimuli might activate HVC or NCM neurons that, in turn, modulate VTA dopaminergic neurons projecting to Area X. This hypothesis aligns with studies suggesting that auditory and motor inputs can influence VTA activity [63-64]. Alternatively, our observed delay in dopamine release may reflect feedback modulation within Area X circuits, further influenced by dopaminergic input from VTA.To clarify these interpretations, future studies could combine optogenetic stimulation of HVC/NCM pathways with simultaneous dopamine measurements in Area X. This would help delineate the causal role of HVC/NCM activity in modulating dopaminergic inputs from VTA to Area X.

From a microelectrode – and specifically material – point of view, this study reinforces the argument that carbon-based materials offer a compelling case for *multi-modality*. The current state-of-the-art research in neural probes predominantly concentrates on either increasing channel counts for electrophysiology, increasing longevity of the implanted microelectrodes, or increasing the voltammetry sensitivity of microelectrodes for neurochemistry applications. As shown in this study, carbon-based electrodes, have significant capability in offering multi-modality with simultaneous and real-time intracortical/synaptic neurotransmitter detection and single-unit and multi-unit electrophysiology recordings. For example, GC microelectrodes have recently been shown to detect dopamine and serotonin at record sensitivities (10 nM) [28,41]. There are still factors that continue to offer obstacles to increased innovations in these carbon materials like non-uniform and sporadic distribution of chemically active functional groups needed to promote surface chemistry for biomolecule adsorption. Recent work in molecular dynamics modeling of the synthesis of such materials may offer some insights on overcoming these shortcomings [60-61].

In conclusion, our study represents a significant advancement in our understanding of the intricate relationship between electrophysiology, neurochemistry, and external stimuli within neural circuits. The innovative three-dimensional probe and advanced analysis techniques presented here have enabled the exploration of these relationships in signficant detail. It is hoped that these findings not only enhance our fundamental comprehension of neural communication but also hold promise for future applications in the diagnosis and treatment of neurological disorders and the development of brain-machine interfaces.

## Acknowledgment

This material is based on research work supported by the Center for Neurotechnology (CNT), a National Science Foundation Engineering Research Center (EEC-1028725), and ONR (Dynamics of Multifunction Brain Network).

## Data Access Statement

Data supporting this study are available upon request.

## Author contributions

N.W.V. designed the experiments, performed the recordings, conducted the analysis, contributed to the probe design, and wrote the manuscript. S.K. proposed the probe concept, wrote the Materials and Methods section, and edited the manuscript.

