## Supplementary material for "Investigating Neurochemistry, Connectivity, and Audio Stimuli Relationship Among Surface and Depth Cortical Neurons": Supplementary_Vahidi etal_Neurochemistry Connectivity AudioStimuliRelations_for bioRxiv_Resub.pdf

Nasim Winchester Vahidi[1†] and Sam Kassegne[2]
[1]Department of Electrical and Computer Engineering, University of California San Diego, La Jolla, California, USA
[2]NanoFAB.SDSU Lab., Department of Mechanical Engineering, College of Engineering, San Diego State University, 5500 Campanile Drive, San Diego, CA, USA

**Section S1. Electrical and Electrochemical Characterizations**
The electrochemical behavior of the microelectrodes was studied in 0.01M PBS (phosphate-buffered saline solution) with pH 7.4 (Sigma Aldrich, USA). Electrochemical impedance spectroscopy (EIS) was used to determine the electrical properties of the probes over a large range of frequencies. using a potentiostat (Reference 600+, Gamry Instruments, USA) connected to a three-electrode electrochemical cell with a platinum wire as a counter electrode and a saturated Ag/AgCl reference electrode. For EIS measurements, 10 mV RMS amplitude sine wave was superimposed on 0 V potential with frequency sweep from 0.1 to $10^5$ Hz. Equivalent circuit modeling of the EIS data was done through Gamry Echem Analyst Vn 7.05 software (Gamry Instruments, USA). Electrical characterization results are shown in **Figure S5**. The mean impedance values at 1 kHz for the epi-intra microelectrodes are 30 kΩ, while that of the voltammetry probes are ~100 kΩ.

**Section S2. Methods for Neural Spike Detection and Sorting**
The following steps were carried out to do neural spike detection and sorting.
**1. Frequency Range:** The selected frequency range (10 Hz to 10 kHz) was designed to encompass both low-frequency local field potentials (LFPs) and high-frequency neural spike activity, providing a broad spectrum of neural activity analysis. This dual focus aligns with the study's objective to investigate the complex relationships between surface and depth neural activities during auditory stimuli responses.
**2. Neural Spike Detection and Sorting:** To ensure the robustness of neural spike detection and sorting. Neural spike data were extracted using band-pass filters in the range of 300 Hz to 4.9 kHz, isolating high-frequency spiking events. Detection thresholds were calculated as a multiple of the signal's standard deviation to minimize noise interference. Spike sorting was performed using principal component analysis (PCA) to differentiate single-unit activity (SUA) from multi-unit activity (MUA), with subsequent manual verification.
**3. Time-Frequency Analysis:** The neural response data were analyzed using spectrograms generated with MATLAB. Specifically. Power spectrograms were created using the Welch method (pwelch function), employing a Hanning window of 128 samples with a 5% overlap. The spectrograms were used to delineate the dominant frequency bands (10–300 Hz for LFPs and 300–10KHz for spikes) to ensure precise interpretation of neural responses and their connectivity.
**4. Addressing "Error Frequency Range" Concern:** The upper frequency limit was capped at 10 kHz to prevent aliasing and retain biologically relevant spiking activity. This was validated during data preprocessing using frequency domain plots to ensure clean separation of LFPs and spikes.

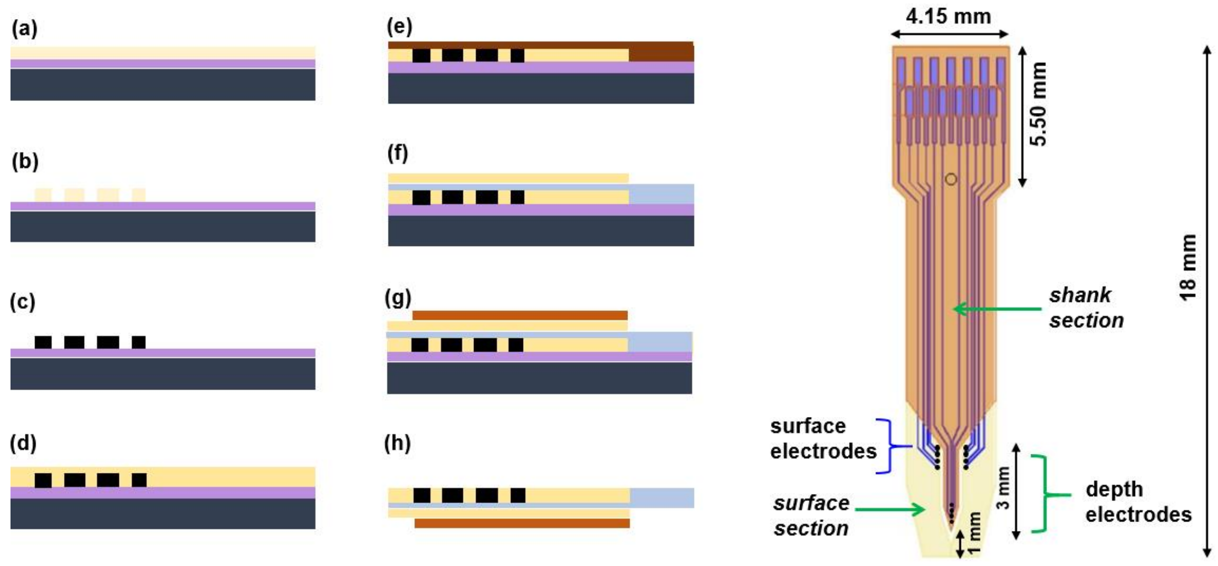


***Figure S1: Epi-Intra probe microfabrication.*** *(a) Spin-coat SU8 negative photoresist on $SiO_2$ substrate. (b) Pattern photoresist through UV exposure. (c) Pyrolyze the patterned resist to obtain GC microelectrodes. (d) Deposit and pattern* base layer of *polyimide (HD 4100) (e) Pattern metal traces through metal lift-off. (f) Add polyimide HD 4100 electrical insulation (g) Reinforce the depth shank with durimide 7520 coating (h) Release the probe from the wafer through buffered hydrofluoric acid (BHF). (i) Schematics of the final Epi-Intra probe showing overall length of 17.5 mm with a depth shank of 3 mm long and 0.5 mm wide.*

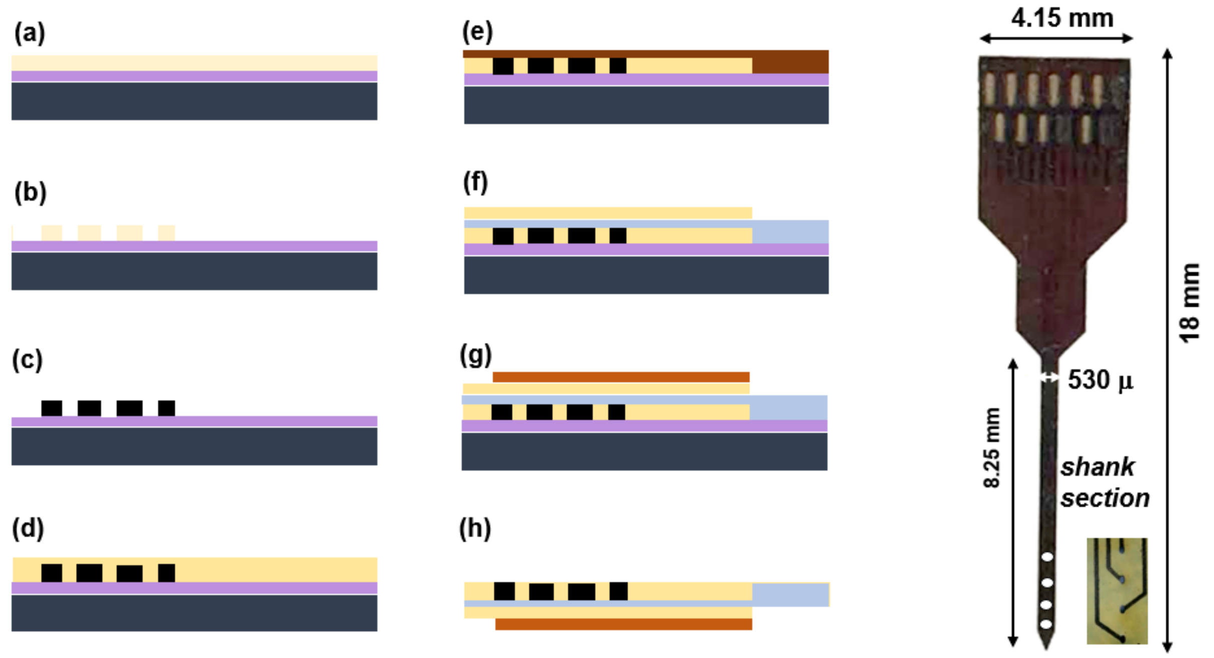


***Figure S2.*** *Shown on left are the microfabrication steps of a penetrating voltammetry probe. The steps are similar as in Figure S1. On the right is shown a 4-channel penetrating neural probes on polymeric substrate (right), with a total shank length of 8.25 mm (and 0.53 mm width) for targeting the rat striatum and four GC microelectrodes of 1500 μm² area, positioned in the striatum, with an inter-electrode distance of 220 μm (inset).*

***(a)*** ***(b)***

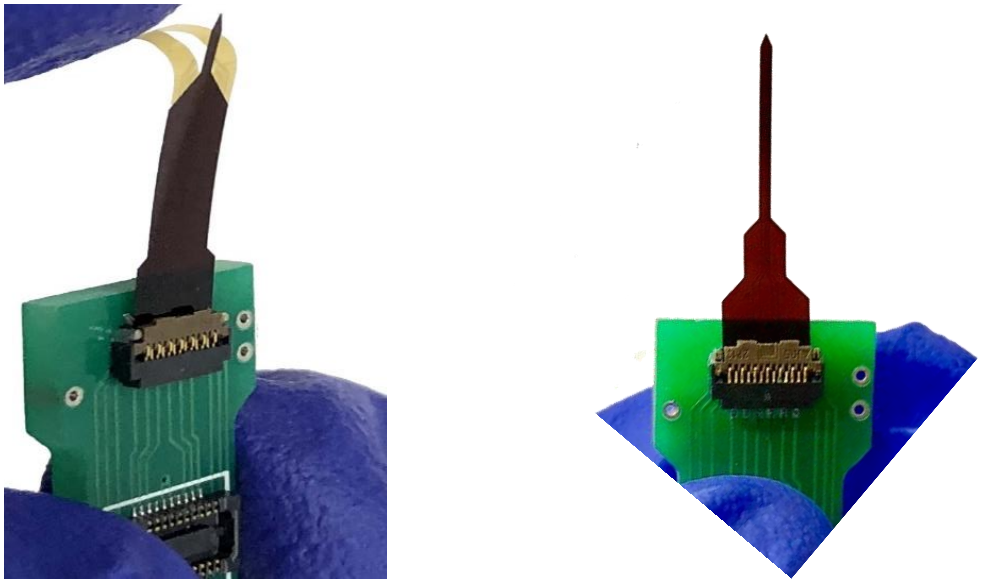

***Figure S3.*** *(a) microfabricated 4.15mm wide and 18.0mm long epi-intra attached to PCB with ZIF connector (b) microfabricated 4-channel voltammetry probe attached to similar PCB.*

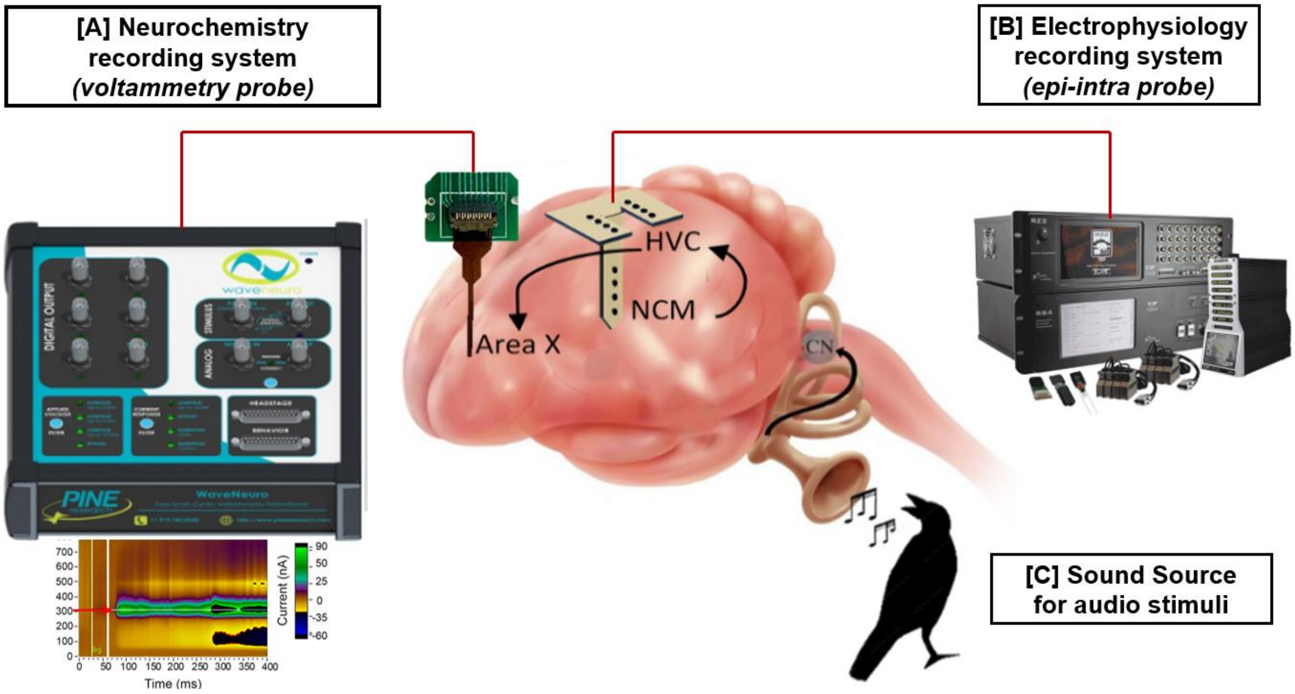


***Figure S4.*** *Schematics of equipment set-up for simultaneous voltammetry and electrophysiology measurements in a songbird. The voltammetry probe was attached to WaveNeuro® FSCV Potentiostat System. The background current is subtracted from the resulting current. Electrophysiological data were acquired using A-M recording systems (Sequim, WA) for 30 min. The neural recording was obtained from NCM auditory area of brain of a European Starling songbird with a sampling rate of 20 kHz. The signals were filtered (300Hz- 5kHz) with gain of 5K. The recorded signals then were sent to a A/D converter (CED Power 1401) with 10X gain to be digitized. Both the epi-intra and voltammetry probes are connected to a PCB that has a*

*ZIF (zero-insertion force) connector. The voltammetry PCBs are then connected to the WaveNeuro FSCV Potentiostat while that of the epi-intra are connected to the A-M recording systems.*

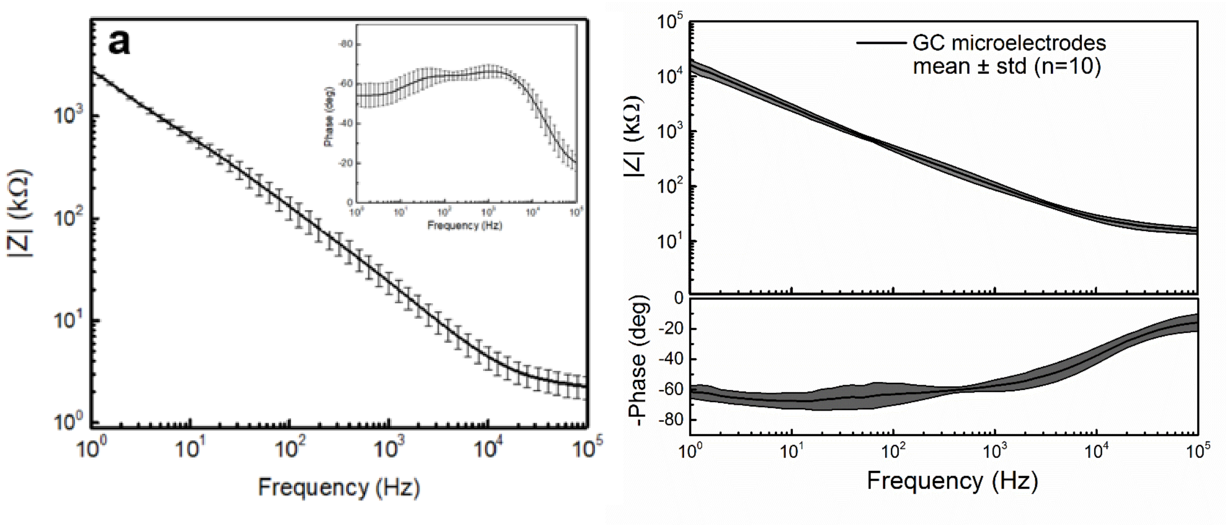


***Figure S5.*** *(a) EIS of Epi-Intra probe with phase shown in the inset, (b) EIS of voltammetry probe.*

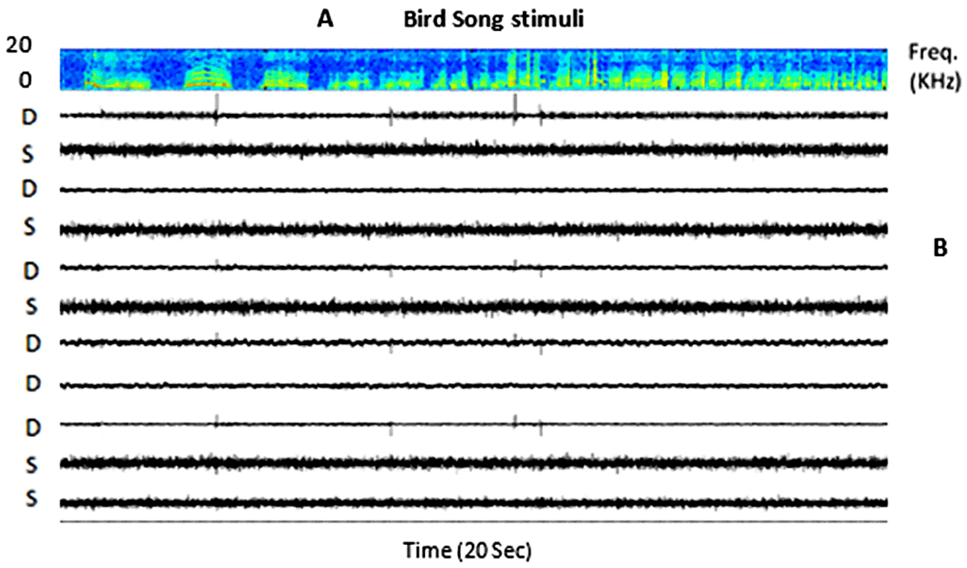


***Figure S6. Example of high-passed data recorded by surface and depth electrodes.*** *(A) Spectrogram of 20-second bird song. The color bar on right indicates power intensity of the song. (B) Response recording by eleven channels.*

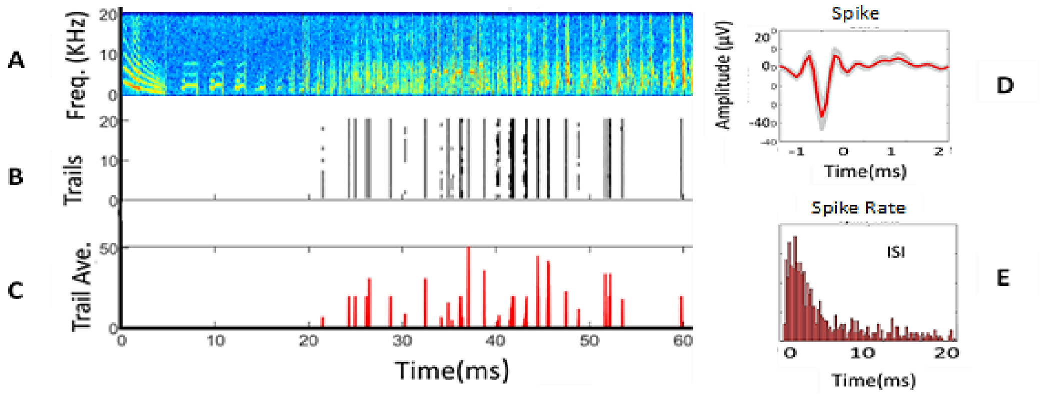


***Figure S7*. *Example a neural spike recorded from NCM*.** *(A) Spectrogram of 60 Sec bird song stimulus. (B Raster plot of spikes for 20 trails of the stimuli (C) Average of the 20 repeated trails. (D) A neural Spike recorded from NCM. The recorded data were analyzed by Mountsort program and spike waveforms is extracted. Red wave is the average of waveforms. (E) Interspike interval of the neural spike.*

**Connectivity Strength dynamics among 11 channels in response to 10 Sec consecutive bird song**

**Connectivity Direction dynamics among 11 channels in response to 10 Sec**

**Correlation dynamics among 11 channels in response to 10 Sec consecutive bird song**

**Neuron-Stimulus Adaptation**

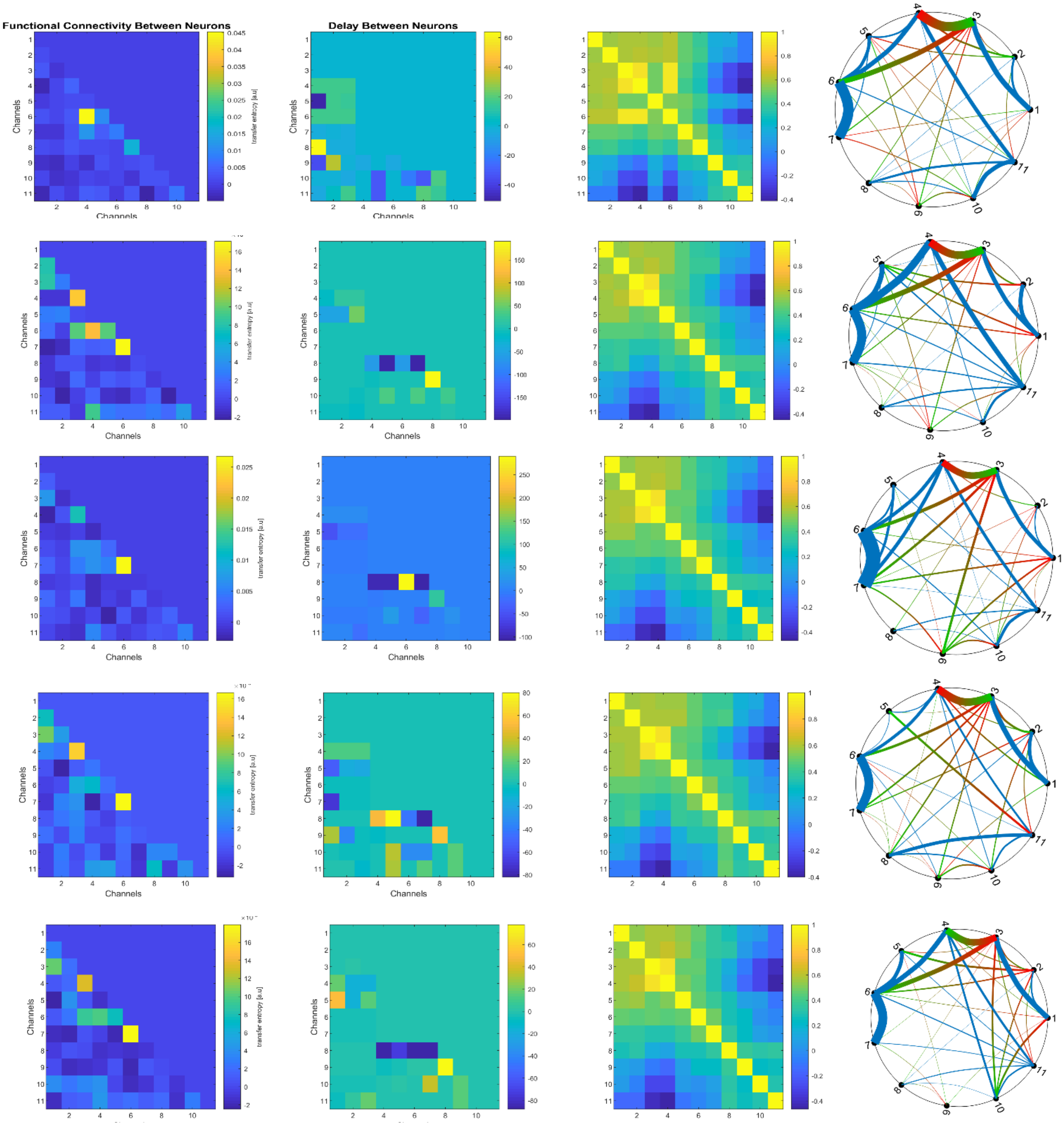

**Connectivity Strength among 11 channels** | **Connectivity Direction among 11 channels** | **Correlation among 11 channels** | **Neuron-Stimulus Adaptation**

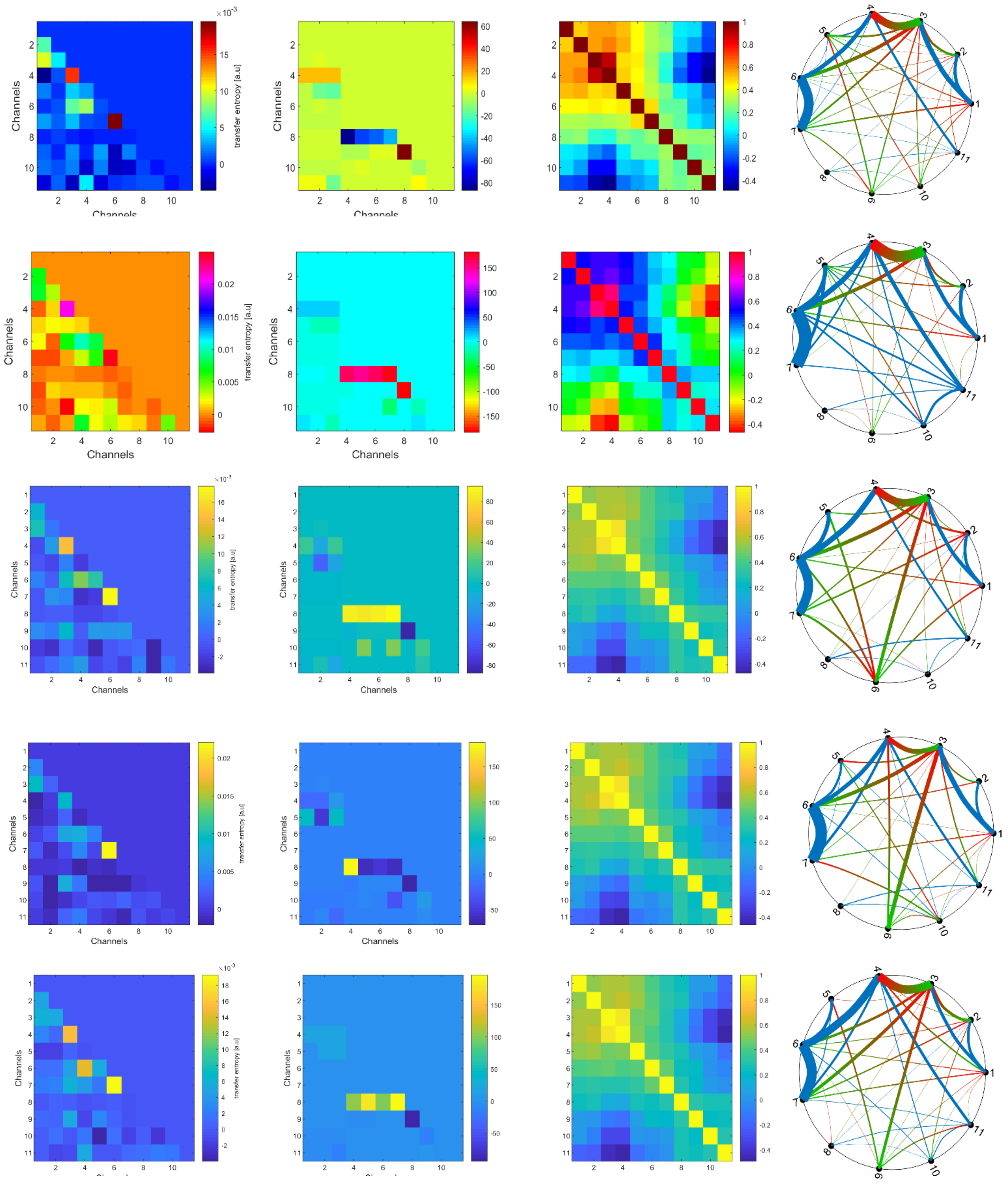

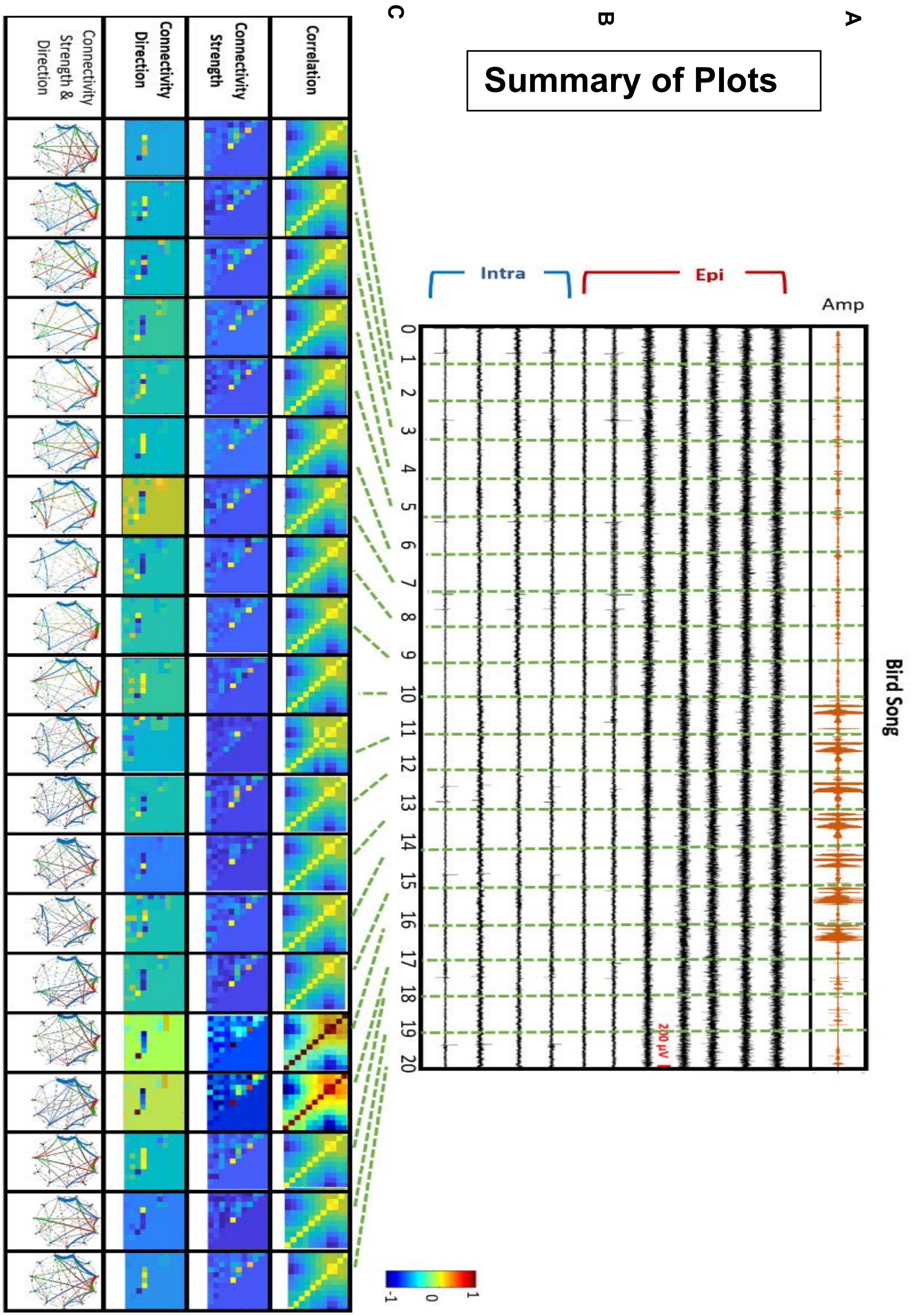


***Figure S8.*** *Extended version of Figure 3.*

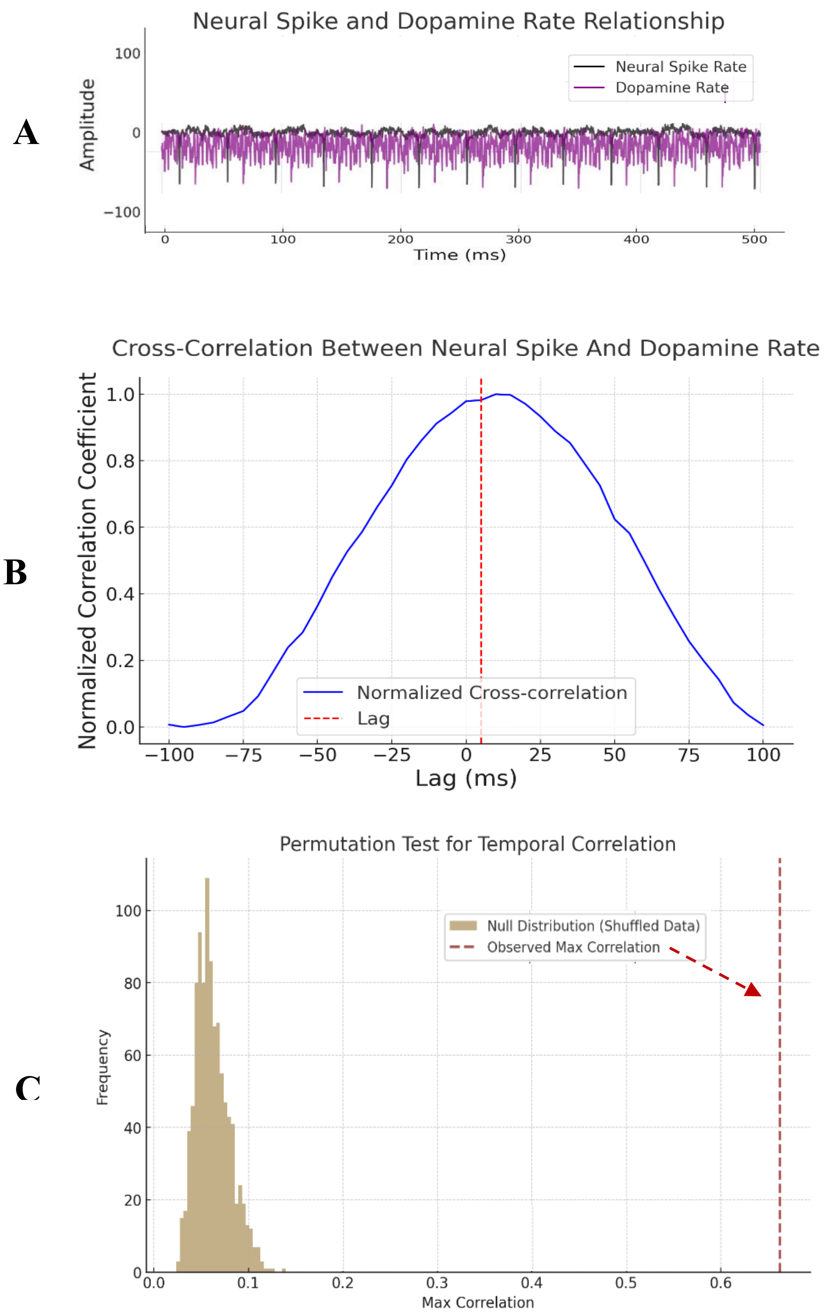


***Figure S9. (A)*** *The plot provides a visualization of the temporal dynamics between NCM neural activity (in black) and dopamine secretion in Area X. (in purple).* ***(B)*** *Time-Lagged Cross-Correlation: To evaluate causality, cross-correlation analysis with time lags was conducted, demonstrating that changes in HVC firing rates precede dopamine current alterations, with a statistically significant peak correlation at the observed delay.* ***(C)*** *Permutation Test**:** The dopamine rate signal was randomly shuffled 1000 times to create a null distribution of maximum correlations. The observed correlation was significantly higher (0.725) than the null distribution (p < 0.01).*
